# Digital Spatial Pathway Mapping Reveals Prognostic Tumor States in Head and Neck Cancer

**DOI:** 10.1101/2025.11.24.689710

**Authors:** Julius Hense, Mina Jamshidi Idaji, Laure Ciernik, Jonas Dippel, Fatma Ersan, Maximilian Knebel, Ada Pusztai, Andrea Sendelhofert, Oliver Buchstab, Stefan Fröhling, Sven Otto, Jochen Hess, Paris Liokatis, Frederick Klauschen, Klaus-Robert Müller, Andreas Mock

## Abstract

Head and neck squamous cell carcinoma (HNSCC) is a morphologically and molecularly heterogeneous disease with limited effectiveness of genotype-informed therapies. Transcriptome-derived estimates of signaling pathway activity carry prognostic and therapeutic potential but remain inaccessible in routine diagnostics due to cost and tissue constraints. Here, we introduce *Digital Spatial Pathway Mapping*, an AI-based computational pathology framework that infers signaling pathway activities directly from routine hematoxylin and eosin (H&E) slides, enabling *in-silico* spatial molecular readouts from standard histology. Models trained on HPV-negative HNSCC from TCGA and externally validated on CPTAC robustly predicted transcriptome-derived activities in cancer-relevant signaling pathways. To achieve spatial interpretability, we applied layer-wise relevance propagation (LRP) to generate heatmaps that highlight positive versus negative evidence for pathway activation. These LRP heatmaps were technically validated by patch-flipping tests and biologically validated against pathway-relevant immunohistochemistry in an independent patient cohort. From these explanations, we derived a *tumor area pathway activity score* (TAPAS) quantifying the spatial fraction of activated tumor regions within a slide. Applied to a retrospective HNSCC cohort of 1,066 slides from 112 resection specimens, TAPAS captured intratumoral heterogeneity and revealed two biologically distinct tumor states - an *oncogenic growth* phenotype with widespread pathway activation and a *pathway quiescent* phenotype associated with higher recurrence risk independent of clinicopathological variables. These findings establish Digital Spatial Pathway Mapping as a scalable, *in-silico* approach to recover systems-level molecular information from standard histopathology, enabling prognostic and mechanistically grounded patient stratification in head and neck cancer.

## 1 Introduction

Head and neck squamous cell carcinoma (HNSCC) represents a heterogeneous group of malignancies arising from the mucosal epithelium of the oral cavity, pharynx, and larynx. The systemic treatment of recurrent and/or metastatic disease remains highly challenging. Immune checkpoint inhibitors benefit only a minority of patients [1], while therapies targeting genetic alterations have achieved little to no clinical success [2, 3]. Current clinical stratification strategies, often limited to mutational profiling from a single tumor block, fail to capture the complexity of intratumoral heterogeneity and microenvironmental context [4, 5].

In this context, transcriptomics is emerging as a powerful frontier in precision oncology [6–8]. Among the most promising readouts are network biology–based estimates of pathway activity [9, 10], the identification of master regulators and cell states [11, 12], as well as deconvolution methods to infer the cellular composition of the tumor microenvironment, increasingly conceptualized in terms of ecotypes [13, 14]. However, the translation of these molecular biomarkers into clinical practice is hampered both by the absence of RNA sequencing in routine diagnostics and by the still incomplete understanding of spatial and temporal intratumoral heterogeneity.

Computational pathology offers a powerful opportunity to bridge the gap between histology and molecular phenotyping [15, 16]. Recent studies have demonstrated that features extracted from routine H&E slides, when integrated with multi-omics data, can accurately predict mutations and transcriptomic profiles across diverse cancer types [17–20]. Critically, this strategy is non-destructive, scalable, and cost-efficient, as it leverages standard diagnostic material without requiring additional assays, making it especially valuable in settings where sequencing is limited or infeasible.

Multiple instance learning (MIL) [21, 22] has emerged as a powerful machine learning paradigm in computational pathology. MIL models aggregate the vast amount of information contained in a histology slide to predict sample- or patient-level endpoints without requiring any spatial annotation for training, a setup known as weakly supervised learning [23–25]. In addition to bulk-level predictions, MIL models can generate spatial attributions within histology slides through heatmaps that highlight regions most relevant to the model prediction. Conventional attention heatmaps [23–25] have been widely used to confirm model focus areas and identify morphologies linked to specific endpoints [26–29]. However, growing evidence shows that attention maps often fail to faithfully capture a model’s decision process [30–35]. Consequently, more advanced explainability methods have been proposed, including additive and SHAP-based MIL approaches [33, 36], as well as xMIL-LRP [34], which combines the explainable multiple instance learning (xMIL) framework with layer-wise relevance propagation (LRP) [16, 35, 37–40]. Unlike attention maps, LRP heatmaps separate positive from negative evidence for a prediction, providing more biologically meaningful spatial insights [34, 41].

From a translational perspective, understanding the spatial distribution of therapeutic targets is essential for drug development and treatment selection. Knowledge of whether a signaling pathway is predominantly active in tumor cells, stromal compartments, or immune infiltrates can help guide targeted therapies.

In this study, we introduce *Digital Spatial Pathway Mapping*, an explainable AI (xAI)-based computational pathology framework that bridges digital histology with systems-level molecular phenotyping. Specifically, we train explainable MIL models on H&E slides to digitally map transcriptome-derived pathway activities and LRP heatmaps that spatially resolve signaling pathways within histological compartments. We demonstrate that these heatmaps faithfully reflect model reasoning, show overlap with pathway-relevant immunohistochemistry and provide interpretable insights into intratumoral heterogeneity. As a quantitative readout of Digital Spatial Pathway Mapping, we define the *tumor area pathway activity score* (TAPAS), which aggregates the fraction of pathway-activated tumor regions in a slide. Applied to a large HPV-negative HNSCC cohort, Digital Spatial Pathway Mapping revealed two biologically distinct tumor states—an *oncogenic growth* phenotype characterized by dependency on oncogenic signaling and potential susceptibility to targeted therapies (e.g., anti-EGFR), and a *pathway quiescent* phenotype largely independent of such signaling and associated with higher recurrence risk.

## 2 Results

### Histology-based prediction of pathway activities on slide-level

We constructed a multiple instance learning (MIL) pipeline to predict signaling pathway activities directly from WSIs of routine H&E-stained tissue sections (**Fig. 1a**). The training dataset comprised 332 cases of HPV-negative HNSCC with matched WSIs and bulk RNA sequencing data from The Cancer Genome Atlas (TCGA). We inferred pathway activities for 14 tumor-relevant signaling pathways from the bulk RNA-seq data using the PROGENy algorithm [9], and thresholded their z-scores at 0.5 to create two target classes: *pathway active* vs. *inactive*. To predict these targets from the WSIs, we extracted patch features with the Virchow2 foundation model [42] and trained Transformer-based MIL models capable of capturing spatial interactions between tissue compartments [25, 34, 43]. We employed 5-fold cross-validation and tested our models on 207 slides from 95 cases of the external HPV-negative validation cohort from CPTAC [44].

**Fig. 1.**
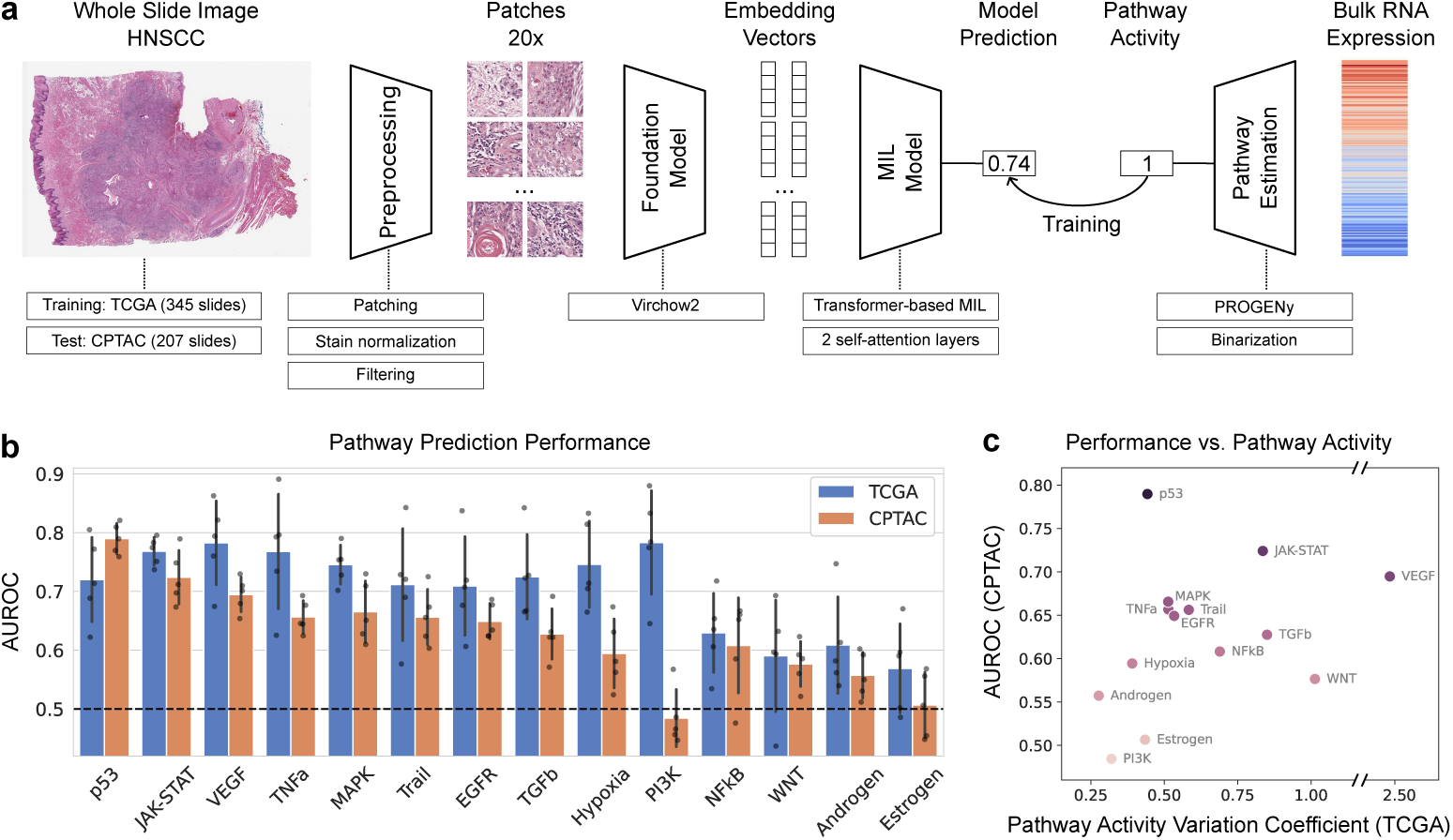
Predicting signaling pathway activity in HNSCC with machine learning. **a)** Multiple instance learning (MIL) pipeline for pathway activity prediction. Models were trained on the TCGA cohort using 5-fold cross-validation. The selected models were evaluated on CPTAC cohort as an independent test dataset. **b)** Predictive performance of the trained models in both the validation set of TCGA (internal validation) and CPTAC (external test) cohorts. The bars show the average AUROC of the five models selected in each fold of the cross-validation (each represented by a dot), and the error bars show the standard deviation across the five folds. **c)** Relationship of the mean AUROC performance on CPTAC external test set vs. the activity variation coefficient (i.e., absolute value of standard deviation divided by mean) of the pathways in TCGA. The dot color corresponds to the mean AUROC on CPTAC (vertical axis).

Many signaling pathways were reliably predicted directly from routine WSIs (**Fig. 1b**). Specifically, our models achieved a mean generalization area under the receiver operating characteristic curve (AUROC) of at least 0.649 on both TCGA and CPTAC for seven of the fourteen pathways. The highest predictive performance was observed for signaling pathways known to play a central role in HNSCC tumorigenesis–such as VEGF, p53, JAK-STAT, TNF*α*, MAPK, EGFR–with mean AUROCs of 0.709-0.783 (TCGA) and 0.649-0.789 (CPTAC). In contrast, pathways less relevant to this tumor type, including Estrogen and Androgen, exhibited comparatively lower performance with mean AUROCs of 0.569-0.609 (TCGA) and 0.507-0.557 (CPTAC). The CPTAC AUROCs were generally attenuated compared to TCGA, suggesting a relevant domain shift across cohorts (*−*10.4% relative performance drop on average; *−*6.9% for top seven CPTAC AUROC pathways).

To explore the variation in predictive performance across pathways, we examined its relation to the heterogeneity of pathway activities in the training cohort. We observed a trend with a moderate positive correlation between the variation of a pathway’s activity across samples in the training data and the generalization performance on the external test data (Spearman’s *ρ* = 0.433, *p*-value = 0.122; **Fig. 1c**). This suggests that greater heterogeneity in pathway activities enabled the model to learn pathway-related morphological markers more effectively, resulting in more robust generalization.

### Digital spatial pathway mapping and validation by immunohistochemistry

After confirming that WSIs can reliably predict slide-level pathway activity labels (i.e., pathway active vs. inactive), we next examined whether such predictions could be spatially resolved to identify the tissue compartments in which pathway activity occurs (e.g., tumor versus non-tumor regions). We employed layer-wise relevance propagation (LRP), which was recently adapted to the MIL framework, for generating interpretable heatmaps [34]. In our application, LRP highlights regions providing positive evidence for the predicted class (pathway active) in contrast to regions contributing negative evidence (pathway inactive), thereby enabling spatially informed insights into pathway activation within histological context (**Fig. 2a**).

**Fig. 2.**
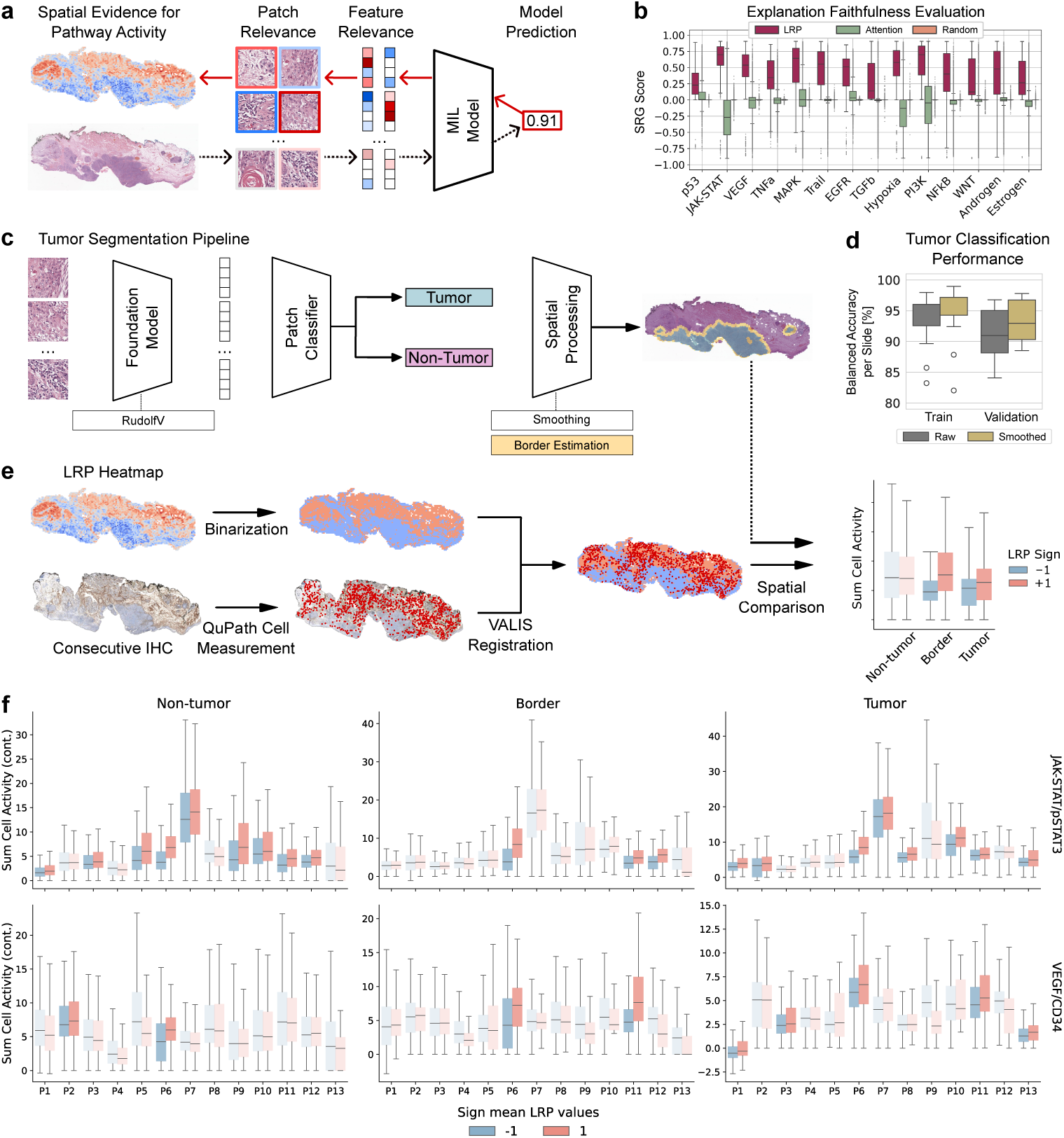
Digital Spatial Pathway Mapping: xAI-based LRP heatmaps and validation with immunohistochemistry. **a)** Schematic of creating LRP heatmaps: each WSI patch is assigned a score reflecting its contribution to the model prediction. Red marks evidence for pathway activity, while blue marks evidence for inactivity. **b)** Faithfulness evaluation using symmetric relevance gain (SRG) scores, applied to 1,066 slides from the LMU cohort. LRP heatmaps show moderate to high alignment with model predictions, while attention-based maps often perform worse than a random baseline. **c)** Overview of the tissue segmentation pipeline, which classifies each patch as tumor, tumor border, or non-tumor. **d)** Classification performance on training and validation data (i.e., 60 and 15 pathologist-annotated slides from the LMU cohort), with a smoothed validation balanced accuracy of 93.47 *±* 3.18 (mean *±* std.) across slides. **e)** Workflow for aligning LRP heatmaps with immunohistochemistry (IHC) and measuring their overlap per tissue compartment. **f)** Spatial overlap analysis results between LRP heatmaps and IHC-derived cell quantifications. Each pair of boxplots (red/blue) shows summed cell activity in positively vs. negatively contributing regions per patient. Solid-colored boxplots indicate statistically significant differences (FDR-adjusted *p <* 0.05).

To technically validate these spatial predictions, we tested whether the heatmaps faithfully represent the decision-making process of the underlying MIL models. Specifically, we conducted a patch-flipping analysis [34, 45] (see Methods): regions with high or low heatmap relevance were systematically removed from a slide, and the resulting changes in the model prediction were measured. Intuitively, a faithful heatmap should cause substantial prediction shifts when highly relevant regions are removed, whereas removing low-relevance regions should leave the predictions largely unchanged. The prediction changes were summarized in the *symmetric relevance gain*(SRG) score [46], where values close to zero indicate random, uninformative heatmaps, and high values indicate faithful alignment with the model’s strategy.

This validation demonstrated that LRP heatmaps accurately reflected the model strategy in the vast majority of cases (**Fig. 2b**). Across pathways, mean SRG scores were significantly above random, exceeding 0.25 in 11 of 14 pathways, confirming strong alignment between heatmap focus areas and model predictions. In contrast, classical attention heatmaps performed no better than random in most cases, underscoring the importance of advanced explainability methods to derive meaningful spatial signals from complex biomarker predictions.

As a biological validation, we assessed whether LRP heatmaps overlap with IHC stains of key pathway-related proteins. Paired IHC–H&E slides were generated for 13 HPV-negative HNSCC samples from our in-house LMU cohort. IHC targeted three pathways with high predictive performance on the slide level: (i) JAK–STAT, using an anti-pSTAT3 antibody; (ii) TNF*α* signaling, using an anti-TNF*α* antibody; and (iii) VEGFA signaling, using an anti-CD34 antibody.

In the context of biology-informed targeted therapy, distinguishing whether pathway activity originates from tumor cells or from immune and stromal compartments is crucial. To address this, we trained a tumor segmentation model that classified each image patch into one of three categories - tumor, tumor border, or non-tumor, achieving a mean balanced accuracy of 93.47% across 15 held-out slides (**Fig. 2c,d**).

To measure the overlap between LRP heatmaps and IHC, cell-level IHC staining intensities were extracted using QuPath [47] after cell segmentation, with IHC slides registered to the H&E WSIs and aggregated at the patch level (see Methods). This yielded a paired dataset of explanation heatmaps and corresponding patch-level cell activation scores for each IHC marker (pSTAT3, TNF*α*, CD34). We then compared staining intensities between regions identified by the heatmaps as positively versus negatively contributing to pathway overactivation, stratified by tissue compartment (**Fig. 2e**). **Fig. S1** illustrates two exemplar slides and their corresponding heatmaps.

As shown in **Fig. 2f**, JAK–STAT heatmaps significantly overlapped with pSTAT3 IHC activation in at least one tissue compartment in 12 of 13 patients, with most of the overlap located in the non-tumor region. Notably, in 7 of these 13 cases, no significant overlap was observed when the analysis was performed without compartmental segmentation (**Fig. S2**), highlighting the importance of fine-grained spatial resolution.

For VEGFA/CD34, significant overlap was detected in 7 of 13 patients when no tissue segmentation is performed (**Fig. S2**). In two cases, overlap was only detectable in the unsegmented analysis, suggesting that signals were distributed across compartments and diluted after stratification. Conversely, in one case, overlap became apparent only after segmentation, indicating that localization effects may be masked in whole-slide analyses.

For TNF*α*, no consistent overlap was observed between explanation heatmaps and IHC staining (**Fig. S2,3**). This suggests that TNF*α* protein abundance, as captured by IHC, may not directly reflect functional pathway activity, underlining the added value of network-based inference from morphology.

In addition, we found that using a foundation model backbone pre-trained on large data cohorts was beneficial for the biological heatmap quality, as we observed weaker overlaps with the IHC stainings when using a foundation model pre-trained on less data (see **Fig. S4-6** and **Supplementary Note 3.2**).

These results demonstrate that Digital Spatial Pathway Mapping can capture spatially resolved, pathway-informed phenotypes directly from H&E slides.

### Digital spatial pathway mapping of intratumoral heterogeneity and association with disease recurrence

Heterogeneity in histology remains underexplored in digital pathology, as most studies analyze only a single representative tumor-containing H&E slide per case, despite surgical specimens typically comprising more than four (or more) tumor-bearing FFPE blocks. To address this limitation, we analyzed a retrospective HPV-negative HNSCC cohort from LMU Munich, Germany (1,066 slides from 112 patients, (**Table S1-2**)), leveraging LRP heatmaps and tumor segmentation to quantify pathway activation within tumor regions of each slide. This yielded the *Tumor Area Pathway Activity Score* (TAPAS), defined as the fraction of tumor patches with positive relevance for pathway activation (see Methods for details). In addition, if more than one (H&E) slide is available per patient, TAPAS can capture intrapatient variation (e.g., within slides from the primary tumor or primary vs. metastatic slides).

Importantly, TAPAS values need to be differentiated from slide-level prediction scores. While the model’s prediction score reflects the pathway activity at the whole-slide level without distinguishing tissue compartments (notably tumor vs. non-tumor regions), TAPAS leverages xAI and quantifies how much the tumor compartment contributes to the model’s decision for the pathway-active class. As a consequence, TAPAS values and slide-level prediction scores only moderately correlate within the LMU cohort (**Fig. S7**).

We computed TAPAS values for all 14 PROGENy pathways and all available (H&E) slides in the LMU cohort. On average, 6.3 primary tumor slides and 3.2 slides of matched lymph node metastases were available per case, providing a comprehensive spatial sampling of tumor heterogeneity. Clustering of TAPAS values revealed two robust and biologically distinct tumor states (**Fig. 3a,b**). The first, designated the *oncogenic growth* state, was characterized by coordinated activation of multiple signaling axes including Hypoxia, MAPK, EGFR, JAK–STAT, TNF*α*, and VEGF. The second, termed the *pathway quiescent* state, showed uniformly low activity across these pathways. The pathway-quiescent state was significantly enriched in slides from lymph node metastases (*p* = 0.003, Fisher’s exact test; **Fig. 3c**), indicating distinct spatial signaling programs between primary and metastatic compartments.

**Fig. 3.**
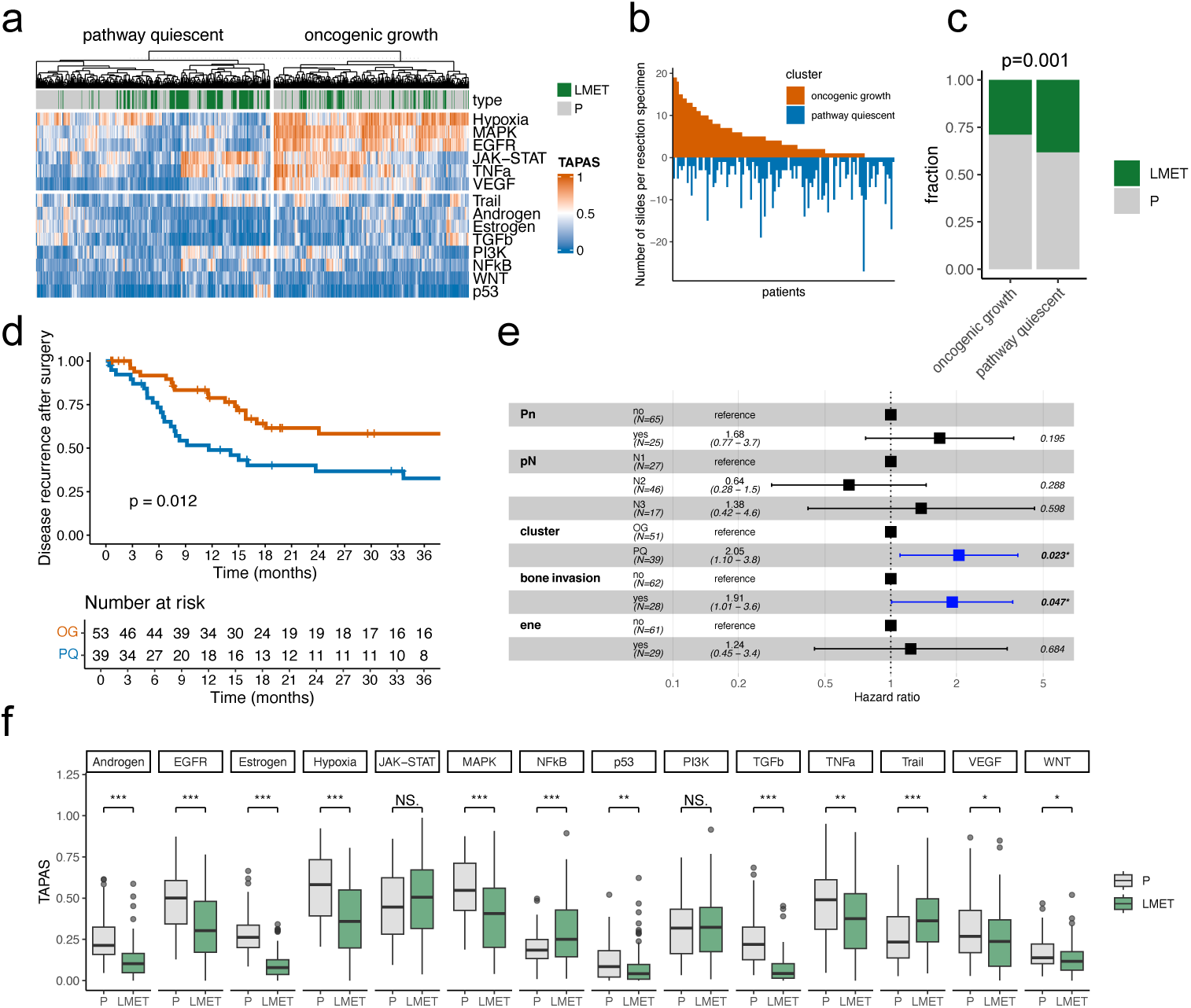
Intratumoral comparison of Digital Spatial Pathway Mapping and survival time modeling. **a)** Heatmap of TAPAS values for all H&E slides in the LMU cohort (1066 slides, 112 patients). Clustering (k-means) revealed a pathway quiescent cluster and a cluster defined by several oncogenic pathway activities (oncogenic growth cluster). **b)** Waterfall plot of the number of slides per resection specimen stratified by clusters. **c)** Fraction of slides classified as oncogenic growth or pathway quiescent cluster for primary (P) and metastatic (LMET) slides. P-value of Fisher’s Exact Test is displayed. **d)** Kaplan-Meier curve of disease-free survival after surgery in the LMU cohort stratified by group. **e)** Forest plot summarizing the multivariate model for all clinicopathological parameters that were significantly associated with disease-free survival in a univariate setting. Significant parameters in the multivariate model are highlighted in blue. **f)** Boxplot of TAPAS values stratified for every pathway and primary vs. lymph node metastasis slides. P-values from t-tests are coded by *p*<*0.05, **p*<*0.01, ***p*<*0.001.

To evaluate clinical relevance, we performed survival time analysis for recurrence-free survival after surgery. Patients classified as pathway quiescent exhibited a significantly increased risk of recurrence (*p* = 0.012, log-rank test; **Fig. 3d**). In univariate analysis, additional parameters associated with recurrence included perineural invasion (Pn), nodal stage (N1–N3), bone invasion, and extranodal extension (ENE) in lymph node metastases (**Table S3**). In multivariate Cox regression, both biological state (pathway quiescent vs. oncogenic growth) and bone invasion remained independent predictors of recurrence (**Fig. 3e**). Interestingly, only TAPAS values and not slide-level prediction scores were independent digital biomarkers in our analayis (**Fig. S8**).

Beyond prognostic associations, the cohort enabled paired comparisons between primary tumors and matched lymph node metastases. When restricting the analysis to tumor regions, TAPAS values differed significantly between primary and metastatic slides for 12 of the 14 PROGENy pathways (**Fig. 3f**). These findings suggest that pathway activation patterns may shift during metastatic progression, potentially reflecting context-dependent dependencies on oncogenic signaling and microenvironmental cues.

Collectively, these results demonstrate that *Digital Spatial Pathway Mapping* enables comprehensive in-silico quantification of signaling pathway activation across all tumor slides, capturing both intra- and intertumoral diversity. The identification of a *pathway quiescent* state associated with poor prognosis highlights the potential of this framework as a spatially resolved digital biomarker for clinically meaningful patient stratification.

## 3 Discussion

This study establishes *Digital Spatial Pathway Mapping* from routine histology as a bridge between tissue morphology and systems-level molecular phenotyping, combining foundation model features, Transformer-based multiple instance learning, and explainable AI. Importantly, we show that the approach generalizes across large public cohorts, yields spatially interpretable readouts, and stratifies patients by recurrence risk.

Our first major contribution is to extend previous work on histology-based prediction of single-gene alterations or expression surrogates [17–20] toward biologically grounded *pathway* activities. Using PROGENy [9], we linked histological features to interpretable signaling axes with direct therapeutic relevance, such as EGFR, JAK–STAT, and VEGF. The successful transfer of models from TCGA to CPTAC underscores the robustness of the approach, while the residual performance drop highlights the persistent challenge of inter-institutional domain shift in digital pathology [48–50].

Second, we contribute to the emerging field of spatially resolved molecular prediction from weakly supervised models. Standard attention heatmaps often fail to faithfully represent model reasoning [30–32], whereas our results demonstrate that using more sophisticated xAI methods like LRP produces technically faithful and biologically plausible explanations [15, 16, 51]. Heatmaps overlapped with pathway-relevant IHC staining for JAK–STAT and VEGF, supporting the biological validity of the spatial signal. Conversely, the absence of concordance for TNF*α* reflects the broader systems-biology insight that pathway activity emerges from network context rather than protein abundance alone [6]. These findings illustrate the potential of xAI to move beyond slide-level predictions toward interpretable maps of pathway activity within tissue compartments.

Third, we systematically quantified intratumoral heterogeneity at scale by computing TAPAS values across all tumor slides, uncovering two robust phenotypes: an “oncogenic growth” state and a “pathway quiescent” state linked to poor outcome. This enabled identification of two robust subgroups: an *oncogenic growth* phenotype with simultaneous activation of multiple pathways, and a *pathway quiescent* phenotype with globally low pathway activity. Strikingly, it was the quiescent state that was associated with significantly worse recurrence-free survival. This counterintuitive observation suggests that diminished canonical pathway activity does not equate to a benign state but may instead capture adverse ecological niches with therapy-resistant cellular quiescence and drug-tolerant persister phenotypes [52–55]. Quiescent tumor populations were shown to evade cytotoxic therapies that preferentially target proliferating cells and can seed relapse after treatment [52, 54]. Notably, the *pathway quiescent* subgroup was enriched in lymph node metastases, suggesting that metastatic niches may favor or select for low-activity states, and that metastatic competence is linked not only to pathway activation but also to plasticity, immune evasion, and stromal interactions.

Methodologically, our integration of pathology foundation models (Virchow2, RudolfV) [42, 56] with Transformer-based MIL [25, 43] represents a robust pipeline for histology-based biomarker prediction. The use of xMIL and LRP [34] overcomes a key limitation of classical attention by explicitly separating evidence for versus against a class, thereby improving interpretability and biological plausibility. Our technical validation through patch flipping and biological validation through IHC registration and quantification establishes a methodological blueprint for rigorous evaluation of digital biomarkers.

From a clinical perspective, image-based pathway prediction offers several advantages. It provides a cost-efficient and non-destructive alternative when RNA-seq is unavailable, limited by tissue constraints, or logistically infeasible [6]. The ability to evaluate all FFPE blocks from a specimen allows more representative assessment of tumor biology than single-sample assays, as highlighted by the prognostic relevance of the *pathway quiescent* subgroup.

While we have performed a retrospective analysis, spatially resolved pathway activities might inform therapeutic hypotheses in individual patients that depend on compartmentalization, such as distinguishing tumor-intrinsic versus stromal or vascular signaling, and may guide rational (combination) therapies targeting EGFR, JAK–STAT, or VEGF pathways [57–59]. Recognition of quiescent, immune-cold states as high-risk could further motivate strategies to reprogram the tumor microenvironment (e.g., TGF*β* blockade or stroma normalization) and to overcome dormancy-associated resistance [54, 60–62].

Several limitations warrant consideration. Our study was restricted to HPV-negative HNSCC, and transferability to HPV-positive disease or other tumor types remains to be tested. The PROGENy pathway panel captures 14 canonical axes but may miss context-specific signaling or master-regulator programs relevant for HNSCC biology. Finally, IHC provides only an imperfect orthogonal reference for validation; future studies should incorporate spatial transcriptomics, proteomics, or functional imaging to triangulate pathway activity with higher resolution. Ultimately, prospective interventional trials will be needed to establish whether histology-derived pathway activities can inform therapeutic decisions and improve patient outcomes.

In conclusion, Digital Spatial Pathway Mapping recovers biologically grounded, spatially resolved pathway activities from routine histology, revealing prognostic tumor states and paving the way for scalable, clinically actionable digital biomarkers in head and neck cancer.

## 4 Methods

### Study cohorts

#### TCGA train cohort

Our training data were extracted from The Cancer Genome Atlas. We downloaded all 472 diagnostic slides (barcode suffix “DX1”-“DX4”) from 450 cases from the TCGA-HNSC project. All slides were taken from the primary tumor (sample type “01”). 53 slides were excluded due to poor quality after pathological review by AM, leaving 419 slides from 405 cases.

The morphological characteristics of head and neck squamous cell carcinoma differ significantly between tumors associated with an HPV infection and those without. To rule out the impact of HPV status on digital biology biomarkers, this study focuses on HPV-negative cases. We obtained information on the HPV status of the patients in the TCGA-HNSC cohort from CBioPortal, PanCancer Atlas 2018. Out of the 405 patients for whom high-quality diagnostic slides were available, 334 were HPV-negative.

We further obtained bulk RNA sequencing samples for the same cohort via recount2 (RSE v.2) [63]. We only considered samples from the primary tumor (sample type “01”) and matched them to the HPV-negative cases with diagnostic slides. For two cases, no bulk RNA-seq data were available. We could assign a unique sample to all other cases, resulting in 345 slides from 332 cases with a matched bulk RNA profile of the primary tumor.

#### CPTAC validation cohort

To construct an independent validation cohort, we used data from The Clinical Proteomic Tumor Analysis Consortium Head and Neck Squamous Cell Carcinoma Collection [44]. We downloaded 390 pathology slides, discarding 175 due to poor quality or missing tumor partitions after review by OB, resulting in 207 slides from 95 cases. All slides were from HPV-negative patients. We obtained RNA sequencing data from Huang et al. [64] and matched them to the remaining cases.

#### LMU cohort

A retrospective cohort of 112 patients diagnosed with HNSCC who underwent surgical resection at the Department of Oral and Maxillofacial Surgery, University Hospital of LMU Munich, between 2011 and 2023 was assembled. A total of 1066 H&E slides were digitized with a high-throughput scanner (Leica Aperio GT 450 DX) at 40× magnification (yielding a resolution of 0.25 µm/pixel). On average, 6.3 primary tumor slides and 3.2 slides of lymph node metastases were digitized per resection specimen. All cases came from the archive of the Institute of Pathology, LMU Munich. An additional 13 cases were used from the archives for the immunohistochemical validations. Here, we created paired H&E and IHC slides for each of the selected three pathways VEGF, JAK-STAT, and TNF*α* (see below). The cohort analysis was approved by the ethics committee of LMU Munich (reference no. 23-0390).

### Data curation

#### Pathway activity inference

The pathway activity for each sample in the TCGA and CPTAC cohorts was inferred using the PROGENy R package [9]. PROGENy leverages a large collection of publicly available perturbation experiments to robustly estimate the activity of 14 tumor-relevant pathways (Androgen, EGFR, Estrogen, Hypoxia, JAK-STAT, MAPK, NFkB, p53, PI3K, TGFb, TNFa, Trail, VEGF, WNT). The input to the algorithm consisted of normalized RNA-seq data (transcript per million; TPM values). For downstream analysis, every pathway was z-scored and binarized using a threshold of 0.5 and categorized as either “active” or “inactive”.

#### Image preprocessing

We tessellated the slides into patches of 340 *×* 340 pixels without overlap at approximately 0.5 microns per pixel (mpp), corresponding to 20x magnification. We identified and excluded background patches via Otsu’s method [65] on slide thumbnails. To eliminate blurry or uninformative patches, we filtered out those with a pixel-based standard deviation below 8.0, without edges according to the Canny edge detection algorithm [66], or with less than two adjacent patches. To mitigate batch effects [50, 67–69], we employed Macenko stain normalization [70]. To maintain consistency across all patches of a slide, we computed color statistics on the slide level by averaging the stain vector estimates of 1,000 randomly sampled patches per slide and applied the resulting normalization operation to all patches of that slide.

We extracted features from all patches using the Virchow2 foundation model [42]. Virchow2 is a publicly available foundation model in histopathology, trained on a diverse dataset of over 3 million histology WSIs. It has demonstrated strong performance as a backbone for both patch-level and slide-level tasks [48, 49, 71, 72]. Moreover, it has been among the most robust models with respect to technical variability (e.g., scanner and staining differences) [50].

### MIL model training

In multiple instance learning (MIL), a model is trained to aggregate the information from multiple instances to predict a single label [21–25]. This weakly supervised learning framework is a popular approach for predicting biomarkers directly from the patches of a whole-slide image [43, 48, 72, 73]. Here, we applied MIL to predict the binarized PROGENy pathway activations (see above). We trained a separate classification model for each of the 14 pathways. As input instances, we used the feature vectors from the Virchow2 foundation model [42] representing the WSI patches.

Recent work indicates that Transformer-based MIL models [25] can outperform the established Attention-based MIL approach [23] in biomarker prediction tasks [43, 48]. Therefore, we employed a Transformer-based MIL architecture comprising an initial multilayer perceptron (MLP) for patch-level feature projection, followed by two multi-head self-attention blocks, using the Nystrom self-attention mechanism [74]. The self-attention mechanism enables the model to capture long-range interactions between the morphological features across the entire WSI. Finally, a linear classification head was applied to the class token to produce a score estimating the probability of activation for the target pathway.

We split the TCGA training dataset at the case level into five equally sized subsets to perform a 5-fold cross-validation. In each fold, one subset was picked for validation, while the remaining four were used for training. We adopted the *train short, test long* strategy: for training, we randomly sampled a fixed number of 2,048 patches per WSI and epoch, while exposing the model to all patches of a slide for prediction. Further training details are presented in the **Supplementary Notes**.

### Explanation heatmaps

#### Heatmap generation

Attention heatmaps are commonly used for the spatial interpretation of a MIL model prediction. They highlight regions that the model focuses on for making its predictions, i.e., features that obtain high attention values inside the model. However, it has been shown that attention heatmaps can be unreliable and particularly unspecific [30–34]. Among other issues, the model may focus on areas in the slide that carry evidence against pathway overactivation and use it to refine the prediction score. As a result, attention heatmaps may misleadingly highlight both pro- and anti-activation regions, combining supportive and contradictory evidence within the same visual interpretation.

Therefore, we utilize xMIL [34], a recently proposed framework for interpreting MIL models. Within xMIL, positive heatmap regions are considered regions of evidence for the predicted class, negative regions against it, and zero regions are neutral. This is in contrast to attention heatmaps that solely highlight focus areas with non-negative attention scores. With this interpretation, we can identify morphologies that support the overactivation of the pathway as those identified as positive.

We utilize the adaptation of Layerwise Relevance Propagation (LRP) for explaining MIL models, which has been shown to yield the most faithful explanations in many settings, particularly in Transformer-based biomarker prediction [34]. In addition to its applicability to MIL models in histopathology, LRP is an xAI method widely used in different domains [16, 35, 37–40, 75].

Starting from the prediction score of a selected class, xMIL-LRP propagates relevance scores through the neurons of all preceding layers according to “propagation rules”. Let *i* be a neuron in layer *l* that receives incoming messages from neurons *j* from subsequent layer *l* + 1, and *q_ij_* the contribution of neuron *i* of layer *l* to relevance 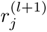. The general propagation rule is defined as

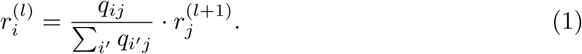

The definition of *q_ij_* depends on the architecture of the respective layer. For attention layers, for example, xMIL-LRP utilizes an attention-specific AH-rule [32]. When reaching the input of the MIL model, the relevance scores of all neurons of a feature vector of a patch are summed up, yielding a single relevance score per patch.

The single scores per patch can be spatially arranged in a heatmap. All positive scores are marked as a red patch, with higher scores obtaining a darker red tone. Similarly, negative scores are shown by blue patches, with lower scores being represented by a darker blue. Zero scores are white. See **Fig. 2a**.

#### Heatmap aggregation

Through the 5-fold cross-validation training procedure, we obtained five models per pathway, resulting in five heatmaps per slide and pathway. Hence, we aggregated heatmaps into a single representative heatmap per slide and pathway:

1. **Mean score**: For each patch, we used the mean of the relevance scores across the five heatmaps as the representative aggregated heatmap score.
2. **Patch exclusion**: We excluded patches with high disagreement among the heatmaps. These patches were masked in the aggregated heatmap visualizations.

a. For each patch, we calculated the standard deviation of relevance scores across all heatmaps and excluded patches with standard deviations above the 0.995 quantile to eliminate highly variable regions.
b. The sign of the heatmap scores is of particular interest as it differentiates between evidence for and against the model prediction. To ensure consistency, we excluded patches where the sign of the mean relevance score differed from the sign in the majority of the heatmaps. We observed that such ambiguous patches typically clustered near the zero boundary.
3. **Outlier clipping**: For visualization purposes, we clipped extreme mean relevance scores to the whisker limits, preserving relative importance while preventing visual domination by outliers.

This approach ensured that the unified heatmaps represented consensus regions of importance while appropriately addressing areas of model disagreement, providing a reliable aggregation of multiple heatmaps for biological interpretation of pathway-associated tissue regions.

### Technical validation of heatmaps using patch flipping

To evaluate how well heatmaps explain the predictions of MIL models, we adopted the concept of faithfulness from explainable AI (xAI) research [45, 46]. Our method, called *patch flipping*, assesses the quality of a heatmap by progressively removing patches from an input slide and observing how the model prediction changes with the modified slide (for details, see [34]). Specifically, the patch flipping procedure works as follows:

1. We use a heatmap to assign an explanation score to each patch in the input slide.
2. We sort the patches based on these scores and divide them into 20 ordered groups. The first group contains the top 5% of patches with the highest scores, the second the patches with 5%-10% highest scores, and so on.
3. We perform two separate experiments, and for each step, we recompute the model prediction on the modified slide.

a. Descending Order: We progressively remove the patches starting with the highest-scoring group (the top 5%) and continue until all patches are dropped.
b. Ascending Order: We do the same, but in reverse, starting with the lowest-scoring group of patches.

By plotting the model prediction against the number of patches removed, we create two perturbation curves, one for the descending order and one for the ascending order. A good heatmap should produce a steep decline in the descending-order curve, since removing the most important patches should quickly change the model prediction. Conversely, it should produce a slow, gradual change in the ascending-order curve, since removing the least important patches should not affect the prediction much. To quantify this intuition, we use the Symmetric Relevance Gain (SRG) metric, which calculates the area between these two curves. A higher SRG value indicates a more faithful and reliable heatmap that reflects the model’s decision-making in a better way. A random heatmap can be expected to result in an SRG score around 0. We conducted the patch-flipping evaluation on all slides from the LMU cohort.

### Tumor segmentation

#### Model training

We required accurate segmentations of tumor regions that allow us to distinguish between spatial pathway signals originating from tumor, tumor border, and non-tumor areas (**Fig. 2c**). To obtain such segmentations in a scalable way, we developed a machine learning model to classify image patches as either tumor or non-tumor tissue. Given that foundation models have demonstrated exceptional performance on comparable classification tasks [56, 71], we trained a lightweight classifier (i.e., a fully connected neural network with a single hidden layer) on top of frozen feature representations from the RudolfV foundation model [56]. We obtained pixel-level tumor region annotations for 75 slides of the LMU cohort from AM. We allocated 60 slides for model training and reserved 15 slides for validation purposes. To generate a distinctive training signal, we labeled patches as tumor if at least 50% of their area fell within the annotated tumor regions. After training, we used the model to classify all patches from the LMU cohort into tumor or non-tumor. Further training and evaluation details are presented in the **Supplementary Note 3.1**.

#### Neighborhood smoothing

To increase the spatial consistency of the tumor predictions, we applied a post-hoc neighborhood smoothing mechanism. For each patch within a slide, we examined the surrounding 3×3 neighborhood of patches and calculated the sum of their continuous prediction scores, which can range from 0 (all patches predicted as non-tumor) to 9 (all patches are predicted as tumor). We applied threshold-based corrections: a sum below 3 resulted in setting the patch prediction to 0, and a sum above 5 resulted in setting the prediction to 1. Otherwise, the prediction remained unchanged. This smoothing process was iteratively repeated until either a stable state with no further prediction changes or a maximum of 100 iterations was reached. We observed that this heuristic approach led to spatially consistent tumor regions and better quantitative prediction performance (see **Fig. 2c**).

#### Border estimation

We applied another postprocessing step to differentiate between distant normal tissue areas and patches in close proximity to the tumor region. Specifically, non-tumor patches within a specified border width of three patches distance from any predicted tumor patch were assigned to a new “border” class. This approach effectively established a spatial buffer zone around tumor regions, allowing us to estimate spatial pathway activity in the tumor microenvironment.

### Biological validation of heatmaps using IHC stains

#### Immunohistochemistry

Immunohistochemical staining for CD34 was performed on a VENTANA Benchmark autostainer (Roche) using the clone QBEnd/10 (Cell Marque, 1:200, 36 minute CC1 pretreatment, incubation 20 minutes, detection system: UltraView).

TNA-alpha (mouse monoclonal antibody, Proteintech, Cat. No. 60291) and pSTAT3 (rabbit polyclonal antibody, Cell Signaling Technology, Cat. No. 9145S) were manually stained using the ImmPRESS Polymer Detection Kits (Vector Laboratories; Anti-Mouse MP-7402, Anti-Rabbit MP-7401).

Antigen retrieval was performed in citrate buffer (Target Retrieval Solution, pH 6, Agilent S2369, 1:10 dilution) by microwave heating at 750 W for two 15-minute cycles, followed by 20 minutes of cooling at room temperature. Endogenous peroxidase activity was blocked with 7.5% H_2_O_2_ for 10 minutes. After blocking with normal serum (ImmPRESS kit, 20 minutes), the primary antibody (TNF-alpha, 1:170; pSTAT3, 1:100 in DAKO Antibody Diluent) was applied for 60 minutes at room temperature. Following two washes in TRIS-Brij buffer, sections were incubated with the respective ImmPRESS Polymer reagent for 30 minutes, developed with DAB+ substrate (Agilent K3467, 3 minutes), rinsed in water, and counterstained.

#### Image registration

We used VALIS [76], a publicly available pipeline to register whole slide images. We registered the IHC slides to the H&E slide using both rigid and non-rigid registrations without cropping. All the overlays of the registered IHC slides with their corresponding H&E slide were manually checked to make sure that the registration has worked properly.

#### Quantification of immunohistochemical stainings

QuPath 0.5.1 was used to quantify the staining intensities determined by immunohistochemistry for the proteins CD34, pSTAT3 and TNF*α* [77]. First, for every protein a staining vector was estimated to optimize the quantification. Next, positive cell detection was performed. Here, optical density sum was used as the detection method. The cell compartment scored was nucleus for pSTAT3 and cell for CD34 and TNF*α*. The intensity parameter threshold was set to 0.05 using a 2.5*µ* pixel size. Cut-offs were set to match the staining intensities to 0.1, 0.2, 0.3 for pSTAT3 and CD34, and 0.2, 0.4, 0.6 for TNFa. Lastly, the cell activation measurements obtained for every slide were exported as a csv file.

#### Analyzing the overlap of IHC activations and heatmaps

Since the H&E slide is taken as the reference slide for the registration procedure, the heatmaps created from model predictions on the H&E slides could be directly overlaid on the newly registered IHC slides. We used the aggregated heatmaps (see above) for this analysis.

For each pair of H&E and IHC slides per patient, we computed the IHC activation for each H&E patch by summing the corresponding IHC signal as measured by QuPath. This yielded two values per patch: a relevance score from the heatmap and an activation score from the IHC slide. We grouped the patches based on the sign of their heatmap scores (positive vs. negative) and conducted a one-sided Mann–Whitney U test to compare the distributions of activation scores. The alternative hypothesis was that patches in negatively scored heatmap regions have lower IHC activation than those in positively scored regions. To account for multiple testing, we applied FDR correction for the tests performed for each pathway.

We first analyzed how the IHC activations within the active and inactive heatmap regions differ for the whole slide tissue. Then, to get a better view of the spatial distribution of the IHC activations, we used the segmentation labels, namely tumor, non-tumor, and tumor border (see above) to overlap the heatmap and IHC stains within each tissue compartment. See **Fig. 2c, d**.

### Survival time modeling

#### Tumor Area Pathway Activation Score (TAPAS)

We combined the segmentation model predictions and the spatial pathway heatmaps to estimate the fraction of the tumor area of a slide that supports pathway over-activation. Specifically, we considered all patches of a slide that were predicted to be tumor by our smoothed segmentation model (see above). Using the aggregated pathway heatmaps, we computed the fraction of these tumor patches that had an aggregated heatmap score greater than zero (and were not excluded during heatmap aggregation due to model disagreement). We refer to this metric as the *Tumor Area Pathway Activity Score* (TAPAS), which we calculated for each slide in the LMU cohort and for each pathway. The metric serves as a computational biomarker of spatial signaling pathway activity inside the tumor.

#### Survival time analysis

Primary endpoint of the Cox proportional hazard modeling was the disease recurrence after surgery. Kaplan-Meier plots were generated with the survminer R package. The ggforestplot R package was used to generate a forest plot of the multivariate model summarizing all clinicopathological parameters that showed a significant association with the endpoint in a univarate setting.

## Declarations

### Data and code availability

Codes for training and evaluating the MIL models and explanation heatmaps are publicly shared at https://github.com/bifold-pathomics/xMIL, and the codes for performing the tumor segmentation, IHC-H&E analyses, as well as patient stratification are shared in the public repository: https://github.com/bifold-pathomics/xMIL-Pathways. The TCGA and CPTAC training and validation cohorts are publicly avail-able via: https://portal.gdc.cancer.gov/. Whole slide images of the LMU validation cohort are not publicly available due to data privacy restrictions.

### Author contributions

Conceptualization and methodology: Ju.H., M.J.I., Jo.H. A.M. Data curation and generation: O.B., A.M., F.E., M.K., P.L., S.O. Creation and evaluation of models and heatmaps: Ju.H., M.J.I., L.C., J.D. Immunohistochemistry: A.S., A.P. Survival analysis: A.M. Analysis of results: Ju.H., M.J.I., L.C., J.D., A.M. Project administration: Ju.H., M.J.I., A.M. Supervision: A.M. Initial draft: Ju.H., M.J.I., L.C., A.M. Manuscript feedback and editing: Ju.H., M.J.I., L.C., J.D., S.F., S.O., Jo.H., F.K., K.-R.M., A.M.

## Supporting information

Hense_et_al_supplements

## Acknowledgements

We thank Aignostics for giving us access to the RudolfV foundation model. The results shown here are in part based upon data generated by the TCGA Research Network: https://www.cancer.gov/tcga.

## References

[1] Ferris, R. L. et al. Nivolumab for recurrent squamous-cell carcinoma of the head and neck. N Engl J Med 375, 1856–1867 (2016).

[2] Keam, B. et al. Personalized biomarker-based umbrella trial for patients with recurrent or metastatic head and neck squamous cell carcinoma: KCSG HN 15-16 TRIUMPH trial. J Clin Oncol 42, 507–517 (2023).

[3] Leemans, C. R., Snijders, P. J. & Brakenhoff, R. H. The molecular landscape of head and neck cancer. Nat Rev Cancer 18, 269–282 (2018).

[4] Gerlinger, M. et al. Intratumor heterogeneity and branched evolution revealed by multiregion sequencing. N Engl J Med 366, 883–892 (2012).

[5] Puram, S. V. et al. Single-cell transcriptomic analysis of primary and metastatic tumor ecosystems in head and neck cancer. Cell 171, 1611–1624.e24 (2017).

[6] Tsimberidou, A. M., Fountzilas, E., Bleris, L. & Kurzrock, R. Transcriptomics and solid tumors: The next frontier in precision cancer medicine. Semin Cancer Biol. 84, 50–59 (2020).

[7] Mock, A., Braun, M., Scholl, C., Fröhling, S. & Erkut, C. Transcriptome profiling for precision cancer medicine using shallow nanopore cDNA sequencing. Sci Rep 13, 2378 (2023).

[8] Mock, A. et al. NCT/DKFZ MASTER handbook of interpreting whole-genome, transcriptome, and methylome data for precision oncology. *npj Precis*. Onc. 7, 109 (2023).

[9] Schubert, M. et al. Perturbation-response genes reveal signaling footprints in cancer gene expression. Nat Commun 9, 20 (2018).

[10] Mock, A. et al. EGFR and PI3K pathway activities might guide drug repurposing in HPV-negative head and neck cancers. Front. Oncol. 11, 678966 (2021).

[11] Alvarez, M. J. et al. Functional characterization of somatic mutations in cancer using network-based inference of protein activity. Nat Genet 48, 838–847 (2016).

[12] Zhou, J. et al. EGFR-mediated local invasiveness and response to cetuximab in head and neck cancer. Mol. Cancer 24, 94 (2025).

[13] Thorsson, V. et al. The immune landscape of cancer. Immunity 48, 812–830.e14 (2018).

[14] Chakravarthy, A. et al. Pan-cancer deconvolution of tumour composition using dna methylation. Nat Commun 12, 1–13 (2021).

[15] Klauschen, F. et al. Toward explainable artificial intelligence for precision pathology. Annu. Rev. Pathol. 19, 541–570 (2024).

[16] Binder, A. et al. Morphological and molecular breast cancer profiling through explainable machine learning. Nat Mach Intell 3, 355–366 (2021).

[17] Kather, J. N. et al. Pan-cancer image-based detection of clinically actionable genetic alterations. Nat Cancer 1, 789–799 (2020).

[18] Fu, Y. et al. Pan-cancer computational histopathology reveals mutations, tumor composition and prognosis. Nat Cancer 1, 800–810 (2020).

[19] Schmauch, B. et al. A deep learning model to predict RNA-seq expression of tumours from whole slide images. Nat Commun 11, 3877 (2020).

[20] Arslan, S. et al. A systematic pan-cancer study on deep learning-based prediction of multi-omic biomarkers from routine pathology images. Commun Med 4, 48 (2024).

[21] Maron, O. & Lozano-Pérez, T. A framework for multiple-instance learning. Advances in Neural Information Processing Systems 10 (1997).

[22] Dietterich, T. G., Lathrop, R. H. & Lozano-Pérez, T. Solving the multiple instance problem with axis-parallel rectangles. Artificial Intelligence 89, 31–71 (1997).

[23] Ilse, M., Tomczak, J. & Welling, M. Attention-based deep multiple instance learning. International Conference on Machine Learning 2127–2136 (2018).

[24] Lu, M. Y. et al. Data-efficient and weakly supervised computational pathology on whole-slide images. Nat Biomed Eng 5, 555–570 (2021).

[25] Shao, Z. et al. Transmil: Transformer based correlated multiple instance learning for whole slide image classification. Advances in Neural Information Processing Systems 34, 2136–2147 (2021).

[26] Lu, M. Y. et al. AI-based pathology predicts origins for cancers of unknown primary. Nature 594, 106–110 (2021).

[27] Lipkova, J. et al. Deep learning-enabled assessment of cardiac allograft rejection from endomyocardial biopsies. Nat Med 28, 575–582 (2022).

[28] Bouzid, K. et al. Enabling large-scale screening of barrett’s esophagus using weakly supervised deep learning in histopathology. Nat Commun 15, 2026 (2024).

[29] Baheti, B. et al. Multimodal explainable artificial intelligence for prognostic stratification of patients with glioblastoma. Modern Pathology 38, 100797 (2025).

[30] Wiegreffe, S. & Pinter, Y. Attention is not not explanation. Proceedings of the 2019 Conference on Empirical Methods in Natural Language Processing and the 9th International Joint Conference on Natural Language Processing (EMNLP-IJCNLP) 11–20 (2019).

[31] Jain, S. & Wallace, B. C. Attention is not Explanation. Proceedings of the 2019 Conference of the North American Chapter of the Association for Computational Linguistics: Human Language Technologies 3543–3556 (2019).

[32] Ali, A. et al. XAI for transformers: Better explanations through conservative propagation. International Conference on Machine Learning 435–451 (2022).

[33] Javed, S. A. et al. Additive MIL: Intrinsically interpretable multiple instance learning for pathology. Advances in Neural Information Processing Systems 35, 20689–20702 (2022).

[34] Hense, J. et al. xMIL: Insightful explanations for multiple instance learning in histopathology. Advances in Neural Information Processing Systems 37, 8300–8328 (2024).

[35] Samek, W., Montavon, G., Lapuschkin, S., Anders, C. J. & Müller, K.-R. Explaining deep neural networks and beyond: A review of methods and applications. Proceedings of the IEEE 109, 247–278 (2021).

[36] Early, J., Evers, C. & Ramchurn, S. Model agnostic interpretability for multiple instance learning. International Conference on Learning Representations (2022).

[37] Bach, S. et al. On pixel-wise explanations for non-linear classifier decisions by layer-wise relevance propagation. PLoS ONE 10 (2015).

[38] Montavon, G., Binder, A., Lapuschkin, S., Samek, W. & Müller, K.-R. Layer-wise relevance propagation: an overview. Explainable AI: interpreting, explaining and visualizing deep learning 193–209 (2019).

[39] Keyl, P. et al. Patient-level proteomic network prediction by explainable artificial intelligence. npj Precis. Onc. 6, 35 (2022).

[40] Keyl, P. et al. Single-cell gene regulatory network prediction by explainable ai. Nucleic Acids Research 51, e20–e20 (2023).

[41] Markey, M. et al. Spatial mapping of gene signatures in hematoxylin and eosin-stained images: A proof of concept for interpretable predictions using additive multiple instance learning. Modern Pathology 38 (2025).

[42] Zimmermann, E., et al. Virchow2: Scaling self-supervised mixed magnification models in pathology. *arXiv preprint arXiv:*2408.00738 (2024).

[43] Wagner, S. J. et al. Transformer-based biomarker prediction from colorectal cancer histology: A large-scale multicentric study. Cancer Cell 41, 1650–1661 (2023).

[44] National Cancer Institute Clinical Proteomic Tumor Analysis Consortium (CPTAC). The clinical proteomic tumor analysis consortium head and neck squamous cell carcinoma collection (cptac-hnscc). 10.7937/k9/tcia.2018.uw45nh81 (2018). Dataset.

[45] Samek, W., Binder, A., Montavon, G., Lapuschkin, S. & Müller, K.-R. Evaluating the visualization of what a deep neural network has learned. IEEE Transactions on Neural Networks and Learning Systems 28, 2660–2673 (2016).

[46] Bluecher, S., Vielhaben, J. & Strodthoff, N. Decoupling pixel flipping and occlusion strategy for consistent XAI benchmarks. Transactions on Machine Learning Research (2024).

[47] Bankhead, P., et al. Qupath: Open source software for digital pathology image analysis. Scientific Reports 16878 (2017).

[48] Neidlinger, P. et al. Benchmarking foundation models as feature extractors for weakly supervised computational pathology. Nat Biomed Eng 1–11 (2025).

[49] Jaume, G. et al. HEST-1k: A dataset for spatial transcriptomics and histology image analysis. Advances in Neural Information Processing Systems 37, 53798–53833 (2024).

[50] Kömen, J., et al. Towards robust foundation models for digital pathology. *arXiv preprint arXiv:*2507.17845 (2025).

[51] Keyl, J. et al. Decoding pan-cancer treatment outcomes using multimodal real-world data and explainable artificial intelligence. Nat Cancer 6, 307–322 (2025).

[52] Sharma, S. V. et al. A chromatin-mediated reversible drug-tolerant state in cancer cell subpopulations. Cell 141, 69–80 (2010).

[53] Hata, A. N. et al. Tumor cells can follow distinct evolutionary paths to become resistant to epidermal growth factor receptor inhibition. Nat Med 22, 262–269 (2016).

[54] Sosa, M. S., Bragado, P. & Aguirre-Ghiso, J. A. Mechanisms of disseminated cancer cell dormancy: an awakening field. Nat Rev Cancer 14, 611–622 (2014).

[55] Pham, T. H., Chen, B., Xu, X., Forman, S. J. & Chen, Y. Cellular quiescence: a promising target for cancer therapy. Nat Rev Cancer 21, 736–750 (2021).

[56] Dippel, J., et al. RudolfV: A foundation model by pathologists for pathologists. *arXiv preprint arXiv:*2401.04079 (2024).

[57] Vermorken, J. B. et al. Platinum-based chemotherapy plus cetuximab in head and neck cancer. N Engl J Med 359, 1116–1127 (2008).

[58] Yu, H., Lee, H., Herrmann, A., Buettner, R. & Jove, R. Revisiting stat3 signalling in cancer: New and unexpected biological functions. Nat Rev Cancer 14, 736–746 (2014).

[59] Kyzas, P. A., Cunha, I. W. & Ioannidis, J. P. A. Prognostic significance of vascular endothelial growth factor immunohistochemical expression in head and neck squamous cell carcinoma: A meta-analysis. Clin Cancer Res 11, 1434–1444 (2005).

[60] Mariathasan, S. et al. Tgf attenuates tumour response to pd-l1 blockade by contributing to exclusion of t cells. Nature 554, 544–548 (2018).

[61] Tauriello, D. V. F. et al. Tgf drives immune evasion in genetically reconstituted colon cancer metastasis. Nature 554, 538–543 (2018).

[62] Jain, R. K. Antiangiogenesis strategies revisited: from starving tumors to alleviating hypoxia. Cancer Cell 26, 605–622 (2014).

[63] Collado-Torres, L. et al. Reproducible RNA-seq analysis using recount2. Nat Biotechnol 35, 319–321 (2017).

[64] Huang, C. et al. Proteogenomic insights into the biology and treatment of HPV-negative head and neck squamous cell carcinoma. Cancer Cell 39, 361–379.e16 (2021).

[65] Otsu, N. et al. A threshold selection method from gray-level histograms. Automatica 11, 23–27 (1975).

[66] Canny, J. A computational approach to edge detection. IEEE Transactions on Pattern Analysis and Machine Intelligence 679–698 (1986).

[67] Howard, F. M. et al. The impact of site-specific digital histology signatures on deep learning model accuracy and bias. Nat Commun 12, 4423 (2021).

[68] Kömen, J., Marienwald, H., Dippel, J. & Hense, J. Do histopathological foundation models eliminate batch effects? A comparative study. NeurIPS 2024 Workshop on Advancements In Medical Foundation Models: Explainability, Robustness, Security, and Beyond. (2024).

[69] de Jong, E. D., Marcus, E. & Teuwen, J. Current pathology foundation models are unrobust to medical center differences. *arXiv preprint arXiv:*2501.18055 (2025).

[70] Macenko, M. et al. A method for normalizing histology slides for quantitative analysis. Proceedings of the IEEE International Symposium on Biomedical Imaging: From Nano to Macro (ISBI) (2009).

[71] kaiko.ai, Gatopoulos, I., Känzig, N., Moser, R. & Otálora, S. eva: Evaluation framework for pathology foundation models. Medical Imaging with Deep Learning (2024).

[72] Campanella, G. et al. A clinical benchmark of public self-supervised pathology foundation models. Nat Commun 16, 3640 (2025).

[73] Campanella, G. et al. Real-world deployment of a fine-tuned pathology foundation model for lung cancer biomarker detection. Nat Med 31, 3002–3010 (2025).

[74] Xiong, Y., et al. Nyströmformer: A Nyström-based algorithm for approximating self-attention, Vol. 35, 14138–14148 (2021).

[75] Montavon, G., Samek, W. & Müller, K.-R. Methods for interpreting and understanding deep neural networks. Digital Signal Processing 73, 1–15 (2018).

[76] Gatenbee, C. D. et al. Virtual alignment of pathology image series for multi-gigapixel whole slide images. Nat Commun 14, 4502 (2023).

[77] Bankhead, P. et al. QuPath: Open source software for digital pathology image analysis. Sci Rep 7, 16878 (2017).

