## Supplementary material for "Digital Spatial Pathway Mapping Reveals Prognostic Tumor States in Head and Neck Cancer": Hense_et_al_supplements

[1]Berlin Institute for the Foundations of Learning and Data, Berlin, Germany.
[2]Machine Learning Group, Technische Universität Berlin, Berlin, Germany.
[3]Aignostics, Berlin, Germany.
[4]Department of Oral and Maxillofacial Surgery and Facial Plastic Surgery, University Hospital, LMU Munich, Germany.
[5]Institute of Pathology, Ludwig Maximilian University of Munich, Munich, Germany.
[6]Division of Translational Medical Oncology, DKFZ, Heidelberg, Germany; NCT Heidelberg, Heidelberg, Germany.
[7]German Cancer Consortium (DKTK), Heidelberg, Germany.
[8]Institute of Human Genetics, Heidelberg University, Heidelberg, Germany.
[9]Department of Otorhinolaryngology, Head and Neck Surgery, University Hospital Heidelberg, Heidelberg, Germany.
[10]German Cancer Consortium (DKTK), partner site Munich, a partnership between DKFZ and Ludwig-Maximilians-Universität München (LMU), Germany.
[11]Korea University, Department of Artificial Intelligence, Seoul, Korea.
[12]MPI for Informatics, Saarbrücken, Germany.
[*]Corresponding author.

[†]These authors contributed equally to this work.

### 1 Supplementary Figures

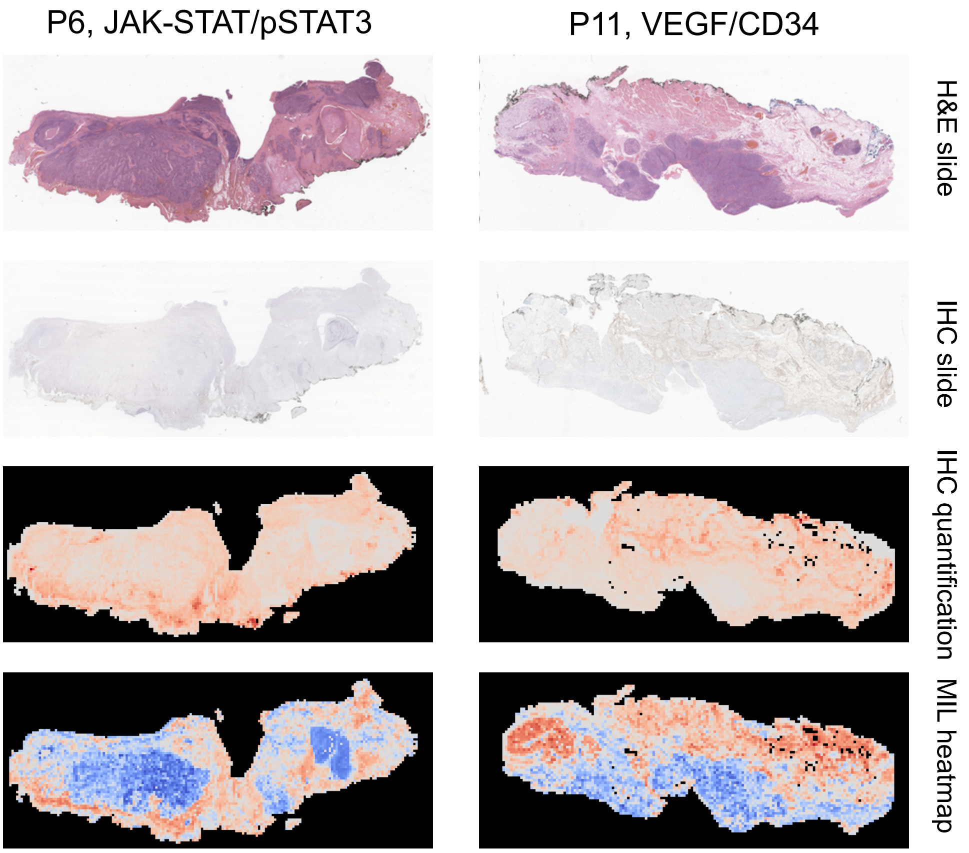


**Fig. S1 Slides and heatmaps for two exemplar patients.** The first two rows show the thumbnails of the H&E and IHC slides (after registration), while row three depicts the heatmaps created from the patch scores of aggregated QuPath measurements within each patch of the H&E slide. The last row illustrates the MIL heatmaps created with LRP.

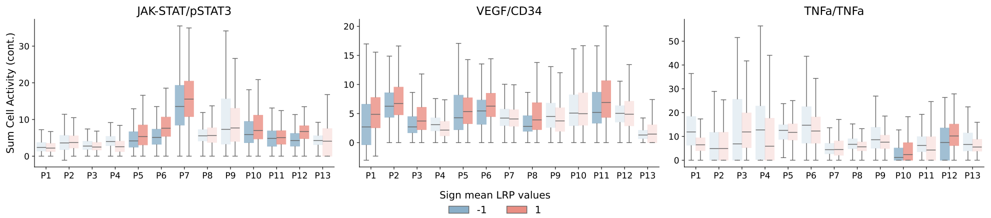


**Fig. S2 Spatial overlap analysis results between heatmaps and IHC-derived cell activations *across all compartments* (without tissue segmentation) with *Virchow2* as the backbone.** Each pair of boxplots (red/blue) shows summed cell activity in positively vs. negatively contributing regions per patient. Solid-colored boxplots indicate statistically significant differences (FDR-adjusted $p < 0.05$).

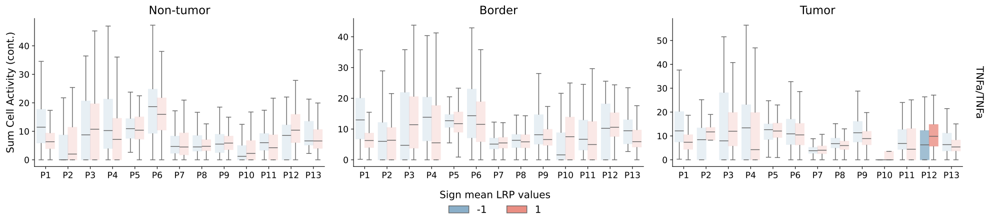


**Fig. S3 Spatial overlap analysis results between heatmaps and IHC-derived cell activations for *TNF$\alpha$* within each tissue compartment with *Virchow2* as the backbone.** Each pair of boxplots (red/blue) shows summed cell activity in positively vs. negatively contributing regions per patient. Solid-colored boxplots indicate statistically significant differences (FDR-adjusted $p < 0.05$).

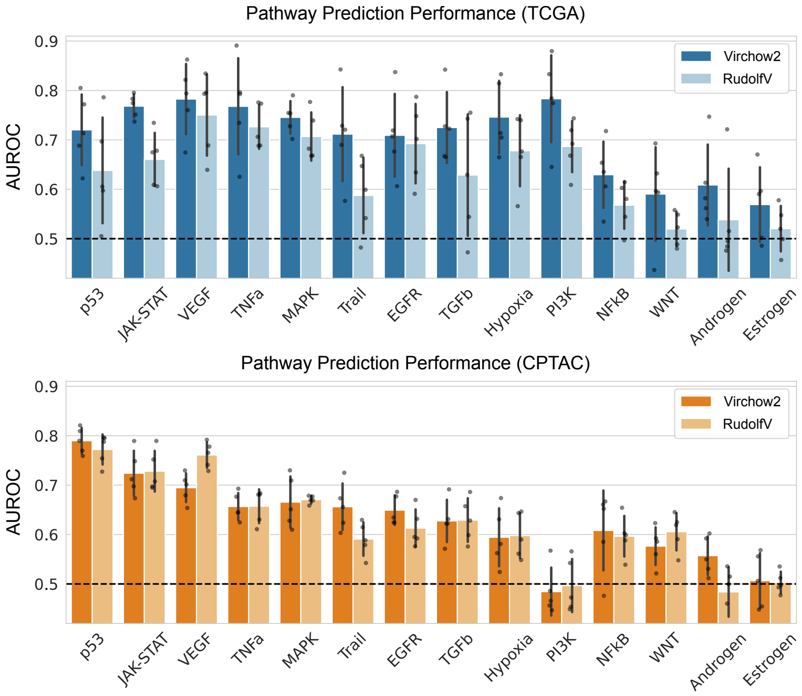


**Fig. S4 Pathway activity prediction using the RudolfV foundation model as backbone.** Predictive performance of the trained models with a *RudolfV* vs. *Virchow2* backbone in both the validation set of TCGA (internal validation) and CPTAC (external test) cohorts. The bars show the average AUROC of the five models selected in each fold of the cross-validation (each represented by a dot), and the error bars show the standard deviation across the five folds.

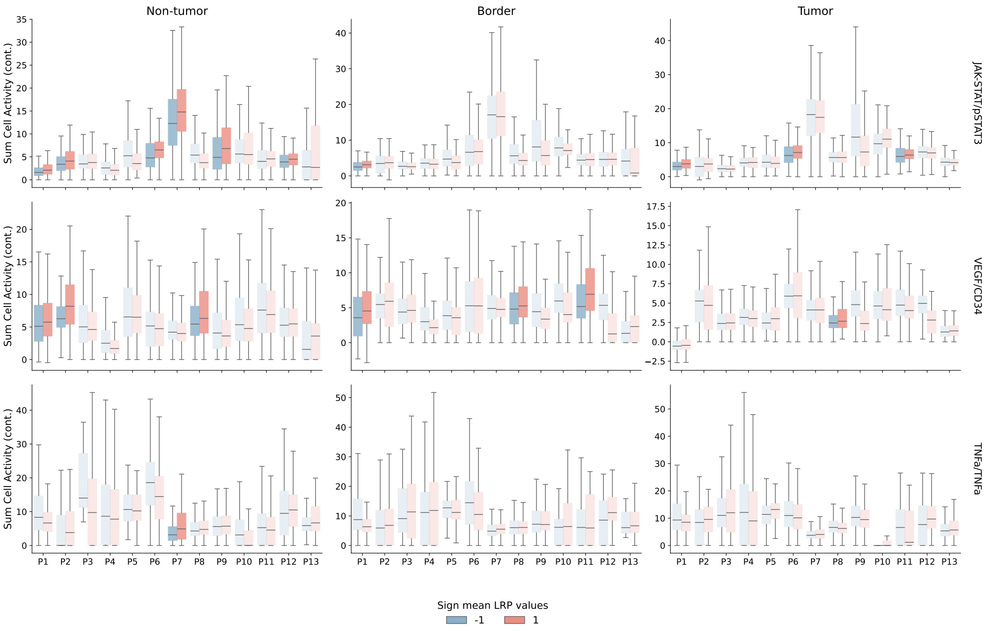


**Fig. S5 Spatial overlap analysis results between heatmaps and IHC-derived cell activations for *all markers* within each tissue compartment with *RudolfV* as the backbone.** Each pair of boxplots (red/blue) shows summed cell activity in positively vs. negatively contributing regions per patient. Solid-colored boxplots indicate statistically significant differences (FDR-adjusted $p < 0.05$).

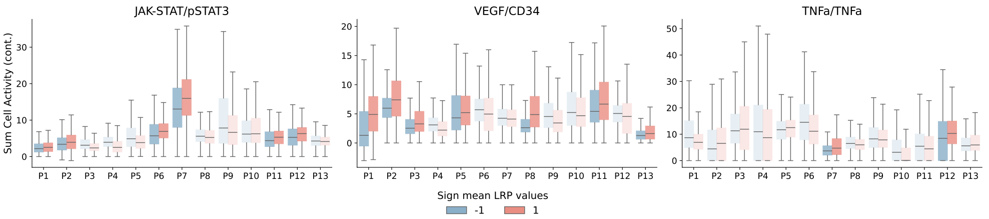


**Fig. S6 Spatial overlap analysis results between heatmaps and IHC-derived cell activations *across all compartments* (without tissue segmentation) with *RudolfV* as the backbone.** Each pair of boxplots (red/blue) shows summed cell activity in positively vs. negatively contributing regions per patient. Solid-colored boxplots indicate statistically significant differences (FDR-adjusted $p < 0.05$).

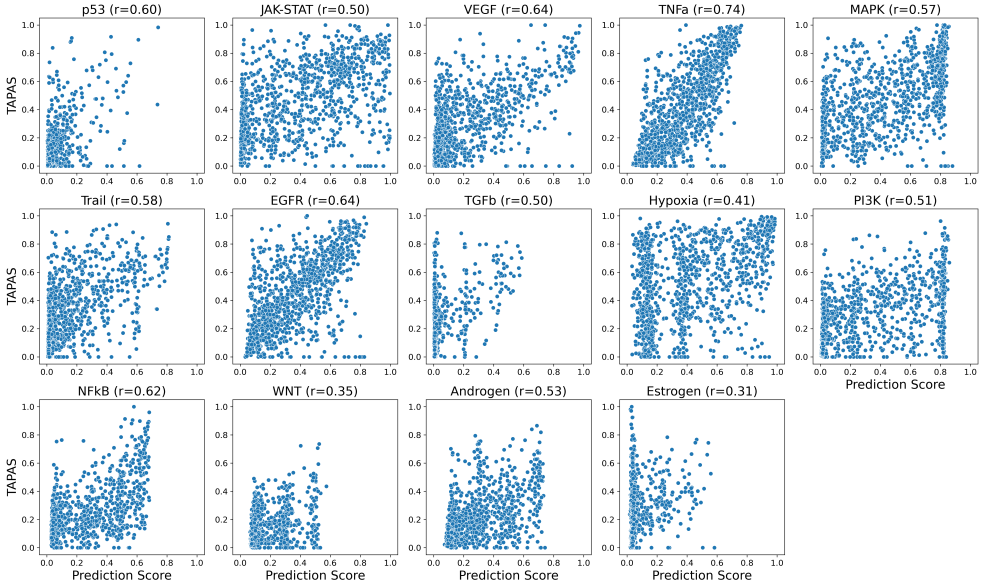


**Fig. S7 Connection between slide-level model prediction and spatial pathway activation.** Softmax prediction score for the pathway activity class averaged over the five MIL models predictions per pathway compared to the spatial tumor area pathway activity score (TAPAS) measured on the LMU cohort. For each pathway, we display the Pearson correlation coefficient r.

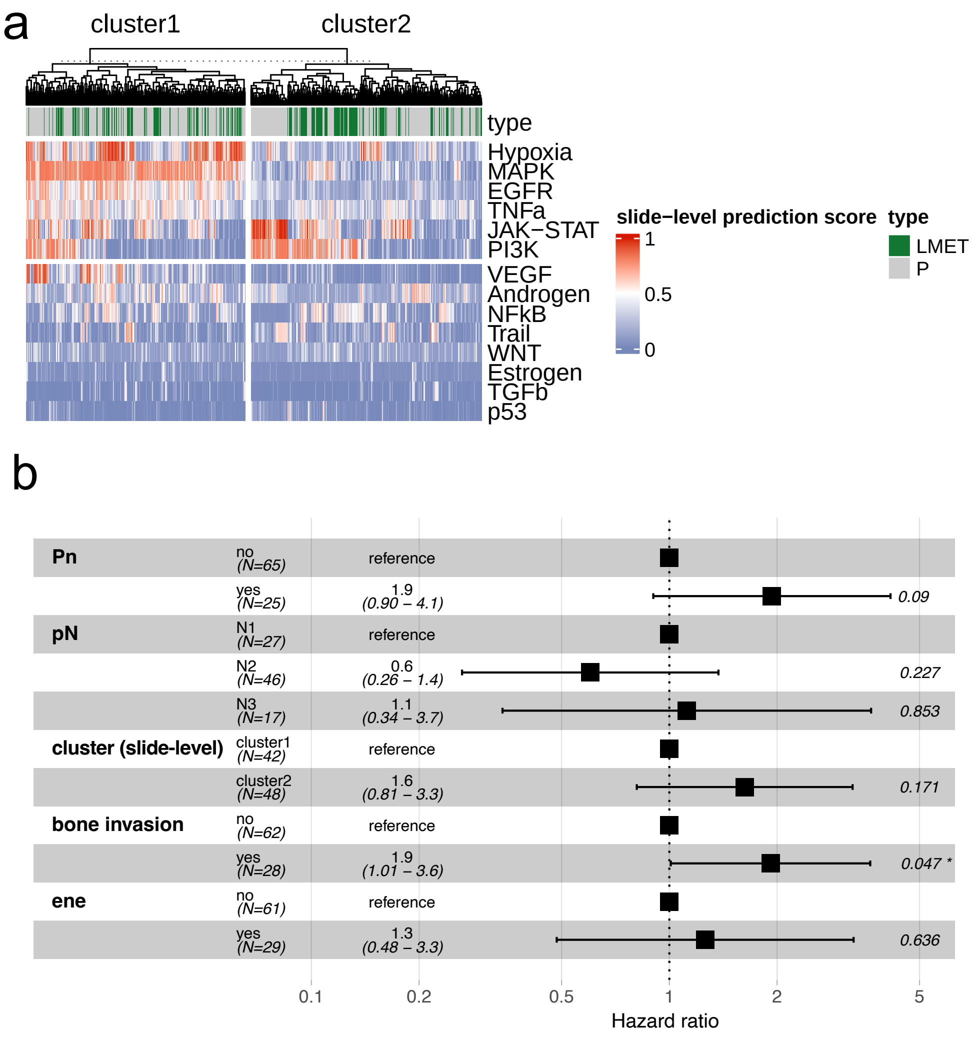


**Fig. S8 Patient stratification in LMU cohort based on slide-level prediction scores. a)** Clustering (k-means) revealed two main clusters. While there is an enrichment of oncogenic pathways in cluster 1, cluster 2 is also characterized by activity in JAK-STAT and PI3K. **b)** Forest plot summarizing the multivariate model for all clinicopathological parameters that were significantly associated with disease-free survival in a univariate setting. P-values are displayed in the top right column.

### 2 Supplementary Tables

**Table S1** Clinicopathologic characteristics of categorial parameters in the LMU cohort (n=112 patients).

| Variable | Level | Count | Percent |
|---|---|---|---|
| L | NA | 2 | 1.8 |
| | no | 67 | 59.8 |
| | yes | 43 | 38.4 |
| Pn | NA | 1 | 0.9 |
| | no | 81 | 72.3 |
| | yes | 30 | 26.8 |
| V | NA | 2 | 1.8 |
| | no | 102 | 91.1 |
| | yes | 8 | 7.1 |
| bone invasion | NA | 1 | 0.9 |
| | no | 77 | 68.8 |
| | yes | 34 | 30.4 |
| ene | NA | 2 | 1.8 |
| | no | 73 | 65.2 |
| | yes | 37 | 33.0 |
| grading | G1 | 6 | 5.4 |
| | G2 | 74 | 66.1 |
| | G3 | 31 | 27.7 |
| | NA | 1 | 0.9 |
| location | Buccal mucose | 9 | 8.0 |
| | Floor of mouth | 30 | 26.8 |
| | Lip | 2 | 1.8 |
| | Mandible | 23 | 20.5 |
| | Maxilla / soft palate | 9 | 8.0 |
| | Retromolar / intramaxillary | 8 | 7.1 |
| | Tongue | 31 | 27.7 |
| pN | N1 | 36 | 32.1 |
| | N2 | 57 | 50.9 |
| | N3 | 19 | 17.0 |
| pT | T1 | 16 | 14.3 |
| | T2 | 34 | 30.4 |
| | T3 | 27 | 24.1 |
| | T4 | 35 | 31.2 |
| radio(-chemotherapy) (yes) | NA | 6 | 5.4 |
| | no | 23 | 20.5 |
| | yes | 83 | 74.1 |
| sex (male) | no | 46 | 41.1 |
| | yes | 66 | 58.9 |

**Table S2** Clinicopathologic characteristics of continous parameters in the LMU cohort (n=112 patients).

| Variable | N | Missing | Median (min–max) | IQR (Q1–Q3) |
|---|---|---|---|---|
| age | 112 | 0 | 67.5 (30.0–93.0) | 59.8–75.2 |
| time from op to recurrence or last follow up(days) | 92 | 20 | 454.0 (11.0–4001.0) | 199.8–1201.0 |
| slides from lymph node metastasis (n) | 112 | 0 | 2.0 (0.0–20.0) | 1.0–4.0 |
| sides from primary tumor (n) | 112 | 0 | 6.0 (0.0–20.0) | 3.0–9.0 |

**Table S3** Univariable Cox regression analysis of clinical and biological variables. Significant p-values (<0.05) are highlighted in bold.

| Variable | Level | HR | 95% CI (low) | 95% CI (high) | p-value | Overall p |
|---|---|---|---|---|---|---|
| Pn | yes | 2.410 | 1.283 | 4.529 | **0.006** | **0.006** |
| pN | N2 | 0.992 | 0.478 | 2.061 | 0.983 | **0.007** |
| pN | N3 | 2.853 | 1.297 | 6.275 | **0.009** | **0.007** |
| cluster | PQ | 2.196 | 1.196 | 4.032 | **0.011** | **0.011** |
| ene | yes | 2.136 | 1.162 | 3.928 | **0.015** | **0.015** |
| bone invasion | ‘bone invasion‘yes | 1.956 | 1.066 | 3.592 | **0.03** | **0.03** |
| V | yes | 3.181 | 0.974 | 10.392 | 0.055 | 0.055 |
| pT | T1 | 0.455 | 0.147 | 1.415 | 0.174 | 0.097 |
| pT | T4 | 1.542 | 0.742 | 3.204 | 0.246 | 0.097 |
| pT | T3 | 0.768 | 0.323 | 1.824 | 0.55 | 0.097 |
| sex (male) | ‘sex (male)‘no | 0.806 | 0.441 | 1.472 | 0.482 | 0.482 |
| age | NA | 1.008 | 0.983 | 1.034 | 0.535 | 0.535 |
| radio(-chemotherapy) (yes) | ‘radio(-chemotherapy) (yes)‘yes | 0.948 | 0.436 | 2.060 | 0.893 | 0.893 |
| L | no | 1.023 | 0.536 | 1.951 | 0.945 | 0.945 |
| location | Floor of mouth | 1.070 | 0.464 | 2.470 | 0.873 | 0.979 |
| location | Retromolar / intramaxillary | 1.168 | 0.326 | 4.190 | 0.811 | 0.979 |
| location | Mandible | 0.968 | 0.401 | 2.337 | 0.942 | 0.979 |
| location | Maxilla / soft palate | 1.577 | 0.548 | 4.545 | 0.399 | 0.979 |
| location | Lip | 0.000 | 0.000 | Inf | 0.997 | 0.979 |
| location | Buccal mucose | 1.408 | 0.447 | 4.438 | 0.559 | 0.979 |
| grading | G3 | 1.058 | 0.543 | 2.061 | 0.869 | 0.987 |
| grading | G1 | 0.000 | 0.000 | Inf | 0.997 | 0.987 |

### 3 Supplementary Notes

#### 3.1 Training details

##### *MIL model training*

We trained all MIL models for 200 epochs with a batch size of five. For each training slide, 2,048 patches were randomly sampled. We applied a dropout scheme to mitigate overfitting, using a dropout rate of 20% after the initial MLP layer and 50% after the self-attention blocks and before the final classification layer. We performed a grid search over learning rates of 0.002 and 0.0002 and training without or with gradient clipping (maximum norm of 1.0). For validation and testing, we always passed the features of all patches of a slide through the MIL model to obtain slide-level predictions. During training, the model checkpoints with the highest validation set AUROCs were saved. For each pathway, we selected the hyper-parameter configuration with the highest mean validation AUROC over the five validation folds, resulting in five separate model checkpoints per pathway. All of these models were then applied to the independent CPTAC test dataset.

##### *Segmentation model training*

We trained a small fully connected neural network with a single hidden layer of 512 dimensions. To address the inherent heterogeneity in tumor sizes across different slides, we employed a hierarchical balanced sampling strategy during training that involved three sequential steps: first, uniformly sampling the class (tumor or non-tumor), then uniformly sampling a slide, and finally uniformly sampling a patch from the selected slide. This approach ensured a sufficient representation of smaller tumors with divergent morphological characteristics that might otherwise be underrepresented. The model was trained for 10 epochs using a batch size of 256, with each epoch sampling an equivalent number of patches to those present in the complete training dataset. After training, we used the model to classify all patches from the LMU cohort into tumor or non-tumor, with each patch receiving a tumor score between 0 and 1 that corresponds to the model's softmax probability for the tumor class. We used 0.5 as a prediction threshold. We measured the model's binary prediction performance on the 15 held-out annotated validation slides by computing the balanced accuracy per slide.

#### 3.2 Pathway prediction results with RudolfV backbone

In a previous attempt, we trained our pathway prediction MIL models using the RudolfV foundation model [1] as a backbone. RudolfV is an earlier foundation model trained on roughly 134K whole-slide images. On performance and robustness benchmarks, Virchow2 [2] outperformed RudolfV, likely due to the much larger pretraining scale [3, 4].

With the RudolfV backbone, most pathways could be predicted as well (see **Fig. S4**). However, the prediction performance on the TCGA validation set was considerably worse than for the Virchow2 model for most pathways (overall mean AUROCs: 0.636 for RudolfV vs. 0.704 for Virchow2), motivating us to update the foundation model backbone to Virchow2 in our pipeline. Interestingly, the generalization performance on the CPTAC cohort only improved for a few pathways (e.g., Trail, EGFR, Androgen) while deteriorating for others, especially VEGF (overall mean AUROCs: 0.622 for RudolfV vs. 0.628 for Virchow2). This may point to comparably strong cross-domain generalization capabilities of RudolfV, as the to-be-expected performance drops were smaller than for Virchow2. Another potential explanation comes from limitations in the CPTAC cohort — due to the worse slide quality than in TCGA, accurate predictions may be more difficult, and the performance potential may be bounded.

Despite the low cross-domain performance drop, the RudolfV-based heatmaps had considerably less overlap with the IHC measurements (see **Fig.** S5). In the tumor regions, statistically significant overlap could only be identified for 3 (JAK-STAT/pSTAT3) and 1 (VEGF/CD34) out of 13 cases, compared to 8 (JAK-STAT/pSTAT3) and 5 (VEGF/CD34) cases with the Virchow2 backbone. Significance in at least one of the three tissue compartments was achieved for 6 (JAK-STAT/pSTAT3) and 4 (VEGF/CD34) cases, compared to 12 (JAK-STAT/pSTAT3) and 6 (VEGF/CD34) cases with Virchow2. That is despite the fact that the number of patients with significant overlap without considering tissue compartments is comparable between RudolfV and Virchow2 (see **Fig.** S6), indicating that the Virchow2-based heatmaps may be more precise. Generally, our results suggest that the explanation heatmap quality can be improved by using larger foundation models that have been pretrained on more WSIs.

### References


[1] Dippel, J. *et al.* RudolfV: A foundation model by pathologists for pathologists. *arXiv preprint arXiv:2401.04079* (2024).

[2] Zimmermann, E. *et al.* Virchow2: Scaling self-supervised mixed magnification models in pathology. *arXiv preprint arXiv:2408.00738* (2024).

[3] Alber, M. *et al.* Atlas: A novel pathology foundation model by Mayo Clinic, Charité, and Aignostics. *arXiv preprint arXiv:2501.05409* (2025).

[4] Kömen, J. *et al.* Towards robust foundation models for digital pathology. *arXiv preprint arXiv:2507.17845* (2025).
